# Evaluating the accuracy of methods for detecting correlated rates of molecular and morphological evolution

**DOI:** 10.1101/2022.07.24.501330

**Authors:** Yasmin Asar, Hervé Sauquet, Simon Y.W. Ho

## Abstract

Determining the link between genomic and phenotypic evolution is a fundamental goal in evolutionary biology. Insights into this link can be gained by using a phylogenetic approach to test for correlations between rates of molecular and morphological evolution. However, there has been persistent uncertainty about the relationship between these rates, partly because conflicting results have been obtained using various methods that have not been examined in detail. We carried out a simulation study to evaluate the performance of five statistical methods for detecting correlated rates of evolution. Our simulations explored the evolution of molecular sequences and morphological characters under a range of conditions. Of the methods tested, Bayesian relaxed-clock estimation of branch rates was able to detect correlated rates of evolution correctly in the largest number of cases. This was followed by correlations of root-to-tip distances, Bayesian model selection, independent sister-pairs contrasts, and likelihood-based model selection. As expected, the power to detect correlated rates increased with the amount of data, both in terms of tree size and number of morphological characters. Likewise, the performance of all five methods improved when there was greater rate variation among lineages. We then applied these methods to a data set from flowering plants and did not find evidence of a correlation in evolutionary rates between genomic data and morphological characters. The results of our study have practical implications for phylogenetic analyses of combined molecular and morphological data sets, and highlight the conditions under which the links between genomic and phenotypic rates of evolution can be evaluated quantitatively.

---

Evolution has generated the great diversity of phenotypic forms across the Tree of Life. However, the genetic mechanisms that underlie changes in phenotype remain incompletely understood (Orr 2001). It is often assumed that there is only a weak link between molecular and morphological change (Simpson 1944; Stanley 1975; Gould and Eldredge 1977; Lee et al. 2013; Halliday et al. 2019), given that the sheer number of mutations accumulating in genomes are likely to include just a small proportion that cause phenotypic changes (Gillespie 1991; Bromham et al. 2002). Furthermore, the proposal of the neutral theory of molecular evolution (Kimura 1968), which describes most mutations as having negligible impact on an organism’s fitness, has bolstered the idea that molecular and morphological evolution are broadly decoupled (Bromham et al. 2002; Davies and Savolainen 2006; Lee et al. 2013; Halliday et al. 2019; Simões et al. 2020). Although genetic drift is believed to be a substantial driver of molecular evolution (Kimura 1968; Ohta 1992), morphological characters, given their importance to an organism’s survival, are often assumed to be under strong selection (Lee and Palci 2015; Ho et al. 2017; Manceau et al. 2020).

Explicit tests of the link between genetic change and phenotypic change have largely focussed on model species, mapping the effect of single genes or small sets of genes (Ashton et al. 2017; Kemble et al. 2019). Recent advances in genomic sequencing have propelled studies of quantitative trait loci, where genomic regions that account for phenotypic trait variation within a species can be identified. However, these studies are time-consuming, costly, and often lack statistical power (Ashton et al. 2017). A more efficient approach that can accommodate hundreds of taxa is to assess the link between rates of molecular and morphological evolution using phylogenetic comparative methods. If there is a correlation between rates of molecular and morphological evolution, then phenotypic traits might be predominantly governed by drift rather than adaptive processes (Halliday et al. 2019), as demonstrated in the evolution of mandibles in rodents (Renaud et al. 2007) and crania in primates (Ackermann and Cheverud 2004). Alternatively, if a species has a high rate of molecular evolution, then it might also have a high rate of morphological evolution simply because it will experience larger numbers of phenotype-altering mutations. However, apparent correlations between molecular and morphological evolutionary rates might be due to both being driven by a third factor. For example, species with small population sizes might experience rapid evolutionary change because of the heightened impacts of drift on both genomic mutations and morphological traits (Combosch et al. 2017).

To date, there have been few explicit tests of the link between molecular and morphological evolutionary rates (Seligmann 2010). Instead, indirect tests have been carried out on ‘living fossils’, or taxa with presumed morphological conservatism across long timescales and high evolutionary distinctiveness (Lidgard and Love 2018; Turner 2019). For instance, the tuatara (*Sphenodon punctatus*) diverged from all other squamates ∼220 million years ago (Ma) and is the only extant member of the order Rhynchocephalia (Herrera-Flores et al. 2017). Despite the inference that the long lineage leading to the tuatara has experienced little morphological evolution (Herrera-Flores et al. 2017; Simões et al. 2022b), the mitochondrial control region of the tuatara has been estimated to evolve at a remarkably high rate (Hay et al. 2008; Subramanian et al. 2009). Earlier work on morphologically conserved horseshoe crabs showed only modest reductions in the rate of mitochondrial evolution compared with scorpions and brine shrimp (Avise et al. 1994). Analysis of *Ginkgo biloba*, the sole surviving species in its order, showed enrichment in duplicated genes and expansion of gene families, bolstering complex chemical defence mechanisms against herbivory (Guan et al. 2016; Šmarda et al. 2016). These results suggest a decoupling of the rates of molecular and morphological evolution in some taxa. In contrast, however, recent transcriptomic analyses of ‘living fossils’ have shown reduced rates of molecular evolution across hundreds of protein-coding genes in *Nautilus* (Combosch et al. 2017; Zhang et al. 2021; Huang et al. 2022; Sanchez et al. 2022), the African coelacanth (Amemiya et al. 2013), the tuatara (Gemmell et al. 2020), and in long-lived sacred lotus (Ming et al. 2013).

Phylogenetic studies of the relationship between morphological and molecular evolutionary rates have produced a mixture of results. An analysis of 13 vertebrate data sets yielded no evidence for an association between rates of molecular and morphological evolution (Bromham et al. 2002), contradicting the results of an earlier study that detected such a correlation in seven of the eight diverse data sets analysed (Omland 1997). Although most studies have tested evolutionary rate correlations in animals, analyses of small data sets from angiosperms (flowering plants) have found weak but positive correlations between rates of molecular and phenotypic evolution (Omland 1997; Barraclough and Savolainen 2001; Davies and Savolainen 2006). These positive correlations have been found across many angiosperm taxa, including: *Sedum* (family Crassulaceae), *Krigia* (family Asteraceae), birch (family Betuleacae), spindle trees (family Celastraceae), Hypoxidaceae, walnuts (family Juglandaceae), *Protea* (family Proteaceae), buckthorns (family Rhamnaceae), and monocots. By gaining insights into the relationship between molecular and morphological evolution in such an extraordinarily species-rich and hyperdiverse group (Onstein 2019), we can achieve a better understanding of the rapid ecological dominance of angiosperms and their evolutionary dynamics (Sauquet and Magallon 2018).

The persistent uncertainty about the relationship between molecular and morphological rates is compounded by the lack of a detailed investigation of the conditions under which a correlation can be detected (Simpson 1944; Seligmann 2010). Without the use of post hoc power analyses (e.g., Davies and Savolainen 2006), it is unclear whether insufficient statistical power in analyses precludes detection of correlated rates or if there is a genuine lack of an association. This is especially pertinent given that previous broad-scale studies examined small sets of genetic markers, commonly ‘housekeeping’ genes (Omland 1997; Barraclough and Savolainen 2001; Davies and Savolainen 2006). These genes might not be well-suited for comparison with morphological rates, since they are under functional constraints and are unlikely to be representative of genome-wide patterns (Bromham et al. 2002). In addition, there remain questions about methodology, such as whether terminal branches of trees inferred with morphological data should be included when testing for correlations between rates of morphological and molecular evolution, given that they might underestimate the amount of evolutionary change (Bromham et al. 2002; Seligmann 2010). This is because autapomorphies (changes that have occurred only in single taxa) are often excluded when collecting morphological characters (Wright and Hillis 2014). Furthermore, previously applied approaches such as root-to-tip distance correlations have been criticized for their time-averaging effect and for the inclusion of non-independent data points, thereby increasing the risk of spurious positive correlations (Felsenstein 1985; Bromham et al. 2002; Rambaut et al. 2016; Barba-Montoya et al. 2021). Thus, it remains unclear whether the mixed results from previous studies have been due to the use of different data sets, insufficient statistical power, or the use of varied methods to test for rate correlations.

Here we aim to uncover the conditions under which correlations between molecular and morphological rates of evolution can be detected. We present a comprehensive simulation study based on parameters from angiosperms, which lends reality to our estimates. We evaluate five approaches for detecting correlated rates of evolution under these conditions, including root-to-tip distance correlations, independent sister-pairs contrasts, likelihood-based model selection, correlations of Bayesian branch rates inferred using relaxed clocks, and Bayesian model selection. Using the insights provided by the simulation study, we test for evolutionary rate correlations between angiosperm floral characters and genomic DNA. Our analyses have implications for understanding the relationship between genotypic and phenotypic change and inform practical recommendations for detecting correlated evolutionary rates.

## Materials and Methods

### Simulations of Molecular and Morphological Evolution

#### Phylogenetic trees and evolutionary rates

We performed simulations using chronograms (branch lengths measured in millions of years, Myr) of three different sizes, based on a 792-species angiosperm tree from Magallón et al. (2015) used by Sauquet et al. (2017). From this tree, we sampled 18, 45, or 111 species to represent diverse angiosperm lineages (Fig. 1a). These three trees had root ages of 139.40 Ma.

**Figure 1.**
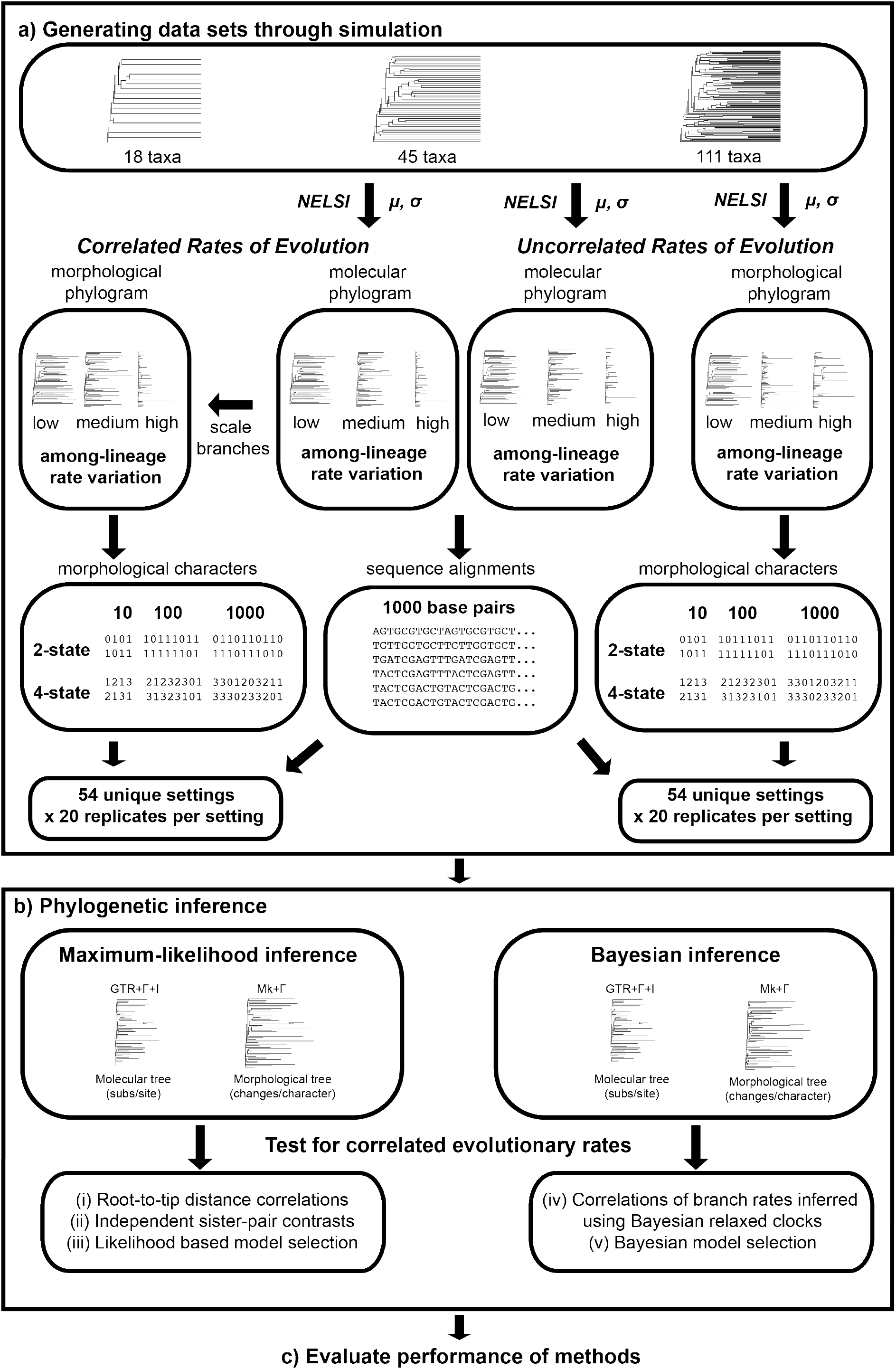
Flowchart of simulation study. a) Generating data sets through simulation. Molecular phylograms are generated using chronograms with three tree sizes (18, 45, or 111 taxa), with a mean rate of evolution (*μ*) that has been inferred from an empirical data set, and standard deviation (*σ*) representing three levels of among-lineage rate variation (0.25, 0.75 or 1.25). The branch lengths of the molecular phylogram are then multiplied by an empirical scaling factor to obtain a morphological phylogram with correlated rates of evolution along each branch. To obtain trees without correlated rates of evolution, morphological phylograms are simply generated using an independent mean rate of evolution (*μ*), but on the same chronogram and with the same standard deviation (*σ*) as the molecular phylogram. Simulations were performed under a total of 54 distinct settings each for the scenario with correlated and uncorrelated rates of evolution, with 20 replicates per setting. b) Phylogenetic inference. Using a fixed tree topology, molecular and morphological branch lengths were inferred using maximum-likelihood and Bayesian analyses. Marginal likelihoods were estimated for the purpose of Bayesian model selection. Five methods were used to test for correlated rates of evolutionary change, (*i*) root-to-tip distance correlations, (*ii*) independent sister-pairs contrasts, (*iii*) likelihood-based model selection, (*iv*) correlations of Bayesian branch rates, and (*v*) Bayesian model selection. c) Evaluate performance of methods. We compared the accuracy and power of the five methods of testing for correlated rates of evolutionary change.

We rescaled the branch lengths of the chronograms to produce phylograms (where branch lengths measured in substitutions/site). To do this, we used the R package *NELSI* (Ho et al. 2015) to generate branch lengths according to an uncorrelated lognormal clock (Drummond et al. 2006). The branch rates had a mean of 9.65×10^−4^ subs/site/Myr, inferred in a previous analysis of two nuclear markers (18S and 26S ribosomal DNA) and three plastid protein-coding genes (*atpB, rbcL*, and *matK*) from 792 angiosperm species (Magallón et al. 2015). We generated branch rates with low, moderate, and high levels of variation, with respective standard deviations of 0.25, 0.75, and 1.25.

We then performed simulations under two scenarios, in which molecular and morphological evolutionary rates were either correlated or uncorrelated. To generate a morphological phylogram with branch rates correlated with those of the molecular phylogram, we scaled the branch lengths of the latter by a factor of 1.90 (Fig. 1a). This scaling factor was based on a mean rate of morphological evolution of 1.83×10^−3^ changes/character/Myr, inferred from 27 floral characters for 792 angiosperm species (Sauquet et al. 2017).

For our simulations with branch rates being uncoupled between molecular and morphological data sets, we simply used *NELSI* to generate independent sets of branch rates and used these to rescale the branch lengths of the chronograms (Fig. 1a). Mean evolutionary rates and standard deviations were as described above for the simulations with correlated branch rates between molecular and morphological data sets.

#### Generating molecular sequence alignments and morphological character matrices

We used Seq-Gen version 1.3.4 (Rambaut and Grass 1997) to simulate the evolution of nucleotide sequences on the phylograms produced by the previous step (Fig. 1a). These simulations produced sequence alignments with lengths of 1000 nucleotides, reflecting typical sizes of nuclear and plastid protein-coding genes. The nucleotide transition rates, frequencies, and gamma shape parameters (degree of among-site rate variation) were based on estimates from Magallón et al. (2015) and are listed in the Supplementary Material.

We generated morphological data matrices consisting of 10, 100, and 1000 characters (Fig. 1a). Matrices with either two-state (binary) or four-state characters were simulated. Character evolution was simulated using the R package *geiger* (Pennell et al. 2014) under the Mk model (Lewis 2001), a generalization of the Jukes-Cantor (1969) model of molecular evolution. The relative rate for each character was drawn randomly from a gamma distribution with a shape parameter of 1.39 inferred via maximum-likelihood analysis of the floral character data set from Schönenberger et al. (2020), comprising 30 binary and multistate morphological characters scored for 792 angiosperm species (see Supplementary Material). This morphological data set was also used in the empirical analyses in this study.

#### Summary of simulation scenarios

The settings described above yielded a total of 54 distinct simulation scenarios. Our simulations included trees of three sizes (18, 45, and 111 taxa). Nucleotide sequences consistently had a length of 1000 nucleotides, but we varied the number of morphological characters (10, 100, and 1000 characters) and the number of possible morphological character states (either two- or four-state characters). We simulated three levels of among-lineage rate heterogeneity with standard deviations of 0.25, 0.75, and 1.25 for the molecular and morphological data sets (Fig. 1a). For each of the 54 simulation settings, we generated 20 replicate data sets. Thus, our simulations produced a total of 1080 pairs of molecular and morphological data sets for each of the ‘correlated’ and ‘uncorrelated’ settings.

### Phylogenetic Inference

#### Maximum-likelihood inference

We inferred phylograms from the simulated molecular and morphological data sets using maximum likelihood in IQ-TREE2 (version ; Bui et al. 2020) (Fig. 1b). In each analysis, the tree topology was constrained to match that used for simulation, with the addition of one outgroup taxon, the gymnosperm *Welwitschia mirabilis*. Analyses of the molecular data used the general-time-reversible model, with gamma rate heterogeneity (four discrete categories) and invariant sites (GTR+Γ+I). Analyses of the morphological data used the time-homogeneous Mk model, with empirical state frequencies and gamma rate heterogeneity with four discrete categories (Mk+Γ) (Supplementary Material). Following maximum-likelihood inference, the gymnosperm outgroup was removed from each tree. The branch lengths from these phylograms were used for root-to-tip distance correlations, independent sister-pairs contrasts, and likelihood-based model selection, which are described in the next section.

#### Bayesian inference

We analysed the molecular and morphological data using Bayesian inference in BEAST2 version 2.6.2 (Bouckaert et al. 2019) (Fig. 1b). In each analysis, the tree topology was constrained to match that used for simulation. Molecular and morphological data were partitioned so that they were assigned distinct substitution models, with the GTR+Γ+I model for the molecular data set and the Mk+ Γ model for the morphological data set. The molecular and morphological data sets were assigned separate uncorrelated lognormal relaxed clocks, with the mean rate of each of these having a uniform prior between 0 and 1. The molecular and morphological data subsets shared the same fixed tree topology, with a birth-death tree prior. We also performed an analysis in which molecular and morphological data shared the same uncorrelated lognormal relaxed clock, to allow comparison between linked and unlinked clock models using Bayesian model selection.

For each Bayesian analysis, the posterior distribution was estimated using Markov chain Monte Carlo (MCMC) sampling. We ran two independent chains, each of 10,000,000 steps, with samples drawn every 1000 steps. The first 10% of samples were discarded as burn-in. After checking for convergence and sufficient sampling (effective sample sizes of parameters greater than 200) using the LogAnalyzer function in BEAST, we combined the samples from the two chains. The tree samples were summarized using TreeAnnotator, part of the BEAST package. We obtained the estimates of branch rates from the summarized trees and used them in our tests of rate correlations, described below.

### Testing for Correlated Rates of Evolution

We evaluated the performance of five methods for testing for correlated evolutionary rates: (*i*) root-to-tip distance correlations, (*ii*) independent sister-pairs contrasts, (*iii*) likelihood-based model selection, (*iv*) correlations of Bayesian branch rates, and (*v*) Bayesian model selection (Fig. 1b). The first three methods were performed on maximum-likelihood trees that were inferred from the simulated data. The last two methods were performed using the results of our Bayesian phylogenetic analyses.

#### (i)Root-to-tip distance correlations

The first method of testing for correlated rates was based on examination of the root-to-tip distances in the maximum-likelihood phylograms. Since all of the tips represent present-day taxa and are separated from the root node by the same amount of time, any differences in the root-to-tip distances reflect differences in evolutionary rates (Omland 1997; Bromham et al. 2002; Arab et al. 2020). For each matched pair of molecular and morphological phylograms, we calculated patristic root-to-tip distances using the distRoot function in the R package *adephylo* (Jombart et al. 2010) and tested for a correlation between the molecular and morphological root-to-tip distances. To address non-independence among the root-to-tip distances, we calculated the *p*-value using a permutation test in the R package *jmuOutlier* (Higgins 2004; Garren 2019), with 20,000 replicates.

#### (ii)Independent sister-pairs contrasts

The second method of testing for correlated rates involved taking phylogenetically independent pairs of taxa (sister pairs) and comparing their relative branch lengths between the molecular and morphological phylograms. We selected sister species that shared a most recent common ancestor to the exclusion of other sister pairs, which avoids the problem of phylogenetic non-independence but reduces the amount of data (Felsenstein 1985; Bromham et al. 2002). We used the R package *diverge* to extract sister pairs for each tree (Anderson and Weir 2022). Lists of sister pairs for each tree size are provided in the Supplementary Material. By definition, the two branches in each sister pair have been evolving for the same amount of time, such that their phylogram length reflects their relative evolutionary rate. We computed the difference between the branch lengths of sister species in the molecular and morphological phylograms inferred using maximum likelihood. We tested for correlations between the molecular and morphological contrasts using the non-parametric Spearman’s rank correlation test, to allow for violations of bivariate normality and homoscedasticity. The contrasts were log-transformed and standardized following standard guidelines by dividing by the square root of the time since divergence between sister species (see Supplementary Materials for details; Garland et al. 1992; Freckleton 2000; Welch and Waxman 2008).

#### (iii).Likelihood-based model selection

The third method of testing for correlated rates involved likelihood-based model selection using information criteria. We analysed the molecular and morphological data using maximum likelihood in IQ-TREE2 and compared models in which the branch lengths were either linked (‘proportionate’) or unlinked between the molecular and morphological trees. These two models reflect correlated and uncorrelated branch rates, respectively, between molecular and morphological data (Duchêne et al. 2020).We compared the models using the corrected Akaike information criterion (AICc) and the Bayesian information criterion (BIC).

#### (iv)Correlations of Bayesian branch rates

The fourth method of testing for correlated rates was based on the inferred branch rates from Bayesian phylogenetic analyses. We tested for correlations between the branch rates inferred using the uncorrelated lognormal relaxed clock for the molecular and morphological data sets. To compute the significance of the correlation, we used Spearman’s rank-order correlation test to avoid violations of bivariate normality and homoscedasticity. For comparison, tests were performed using the mean and median posterior branch rates. The mean posterior rate is commonly reported in Bayesian phylogenetic analyses, but the median rate might be more appropriate because the marginal posterior distributions are skewed.

#### (v)Bayesian model selection

The fifth method of testing for correlated rates involved the use of Bayesian model selection to compare the support for linked or unlinked branch rates between molecular and morphological data. Model selection was performed using Bayes factors, which compare the marginal likelihoods of the two models. The marginal likelihoods of the linked and unlinked relaxed-clock models were estimated using the ‘MS’ Model Selection package in BEAST2. We used generalized-stepping-stone sampling (Baele et al. 2016), with 25 steps and chain lengths of 400,000. The ratio of the marginal likelihoods was used to compute the Bayes factor, which was then interpreted using the guidelines of Kass and Raftery (1995).

#### Evaluation of performance

We used two approaches to evaluate the performance of the five methods of testing for rate correlations. First, we evaluated the methods by their power, which is the ability to detect positive correlations under the largest number of scenarios. Second, we examined the accuracy of the methods, regarded here as the ability to detect positive correlations without false positives. We do not attempt to evaluate the ability of the methods to infer the correct *degree* of rate correlation, i.e., the correlation coefficient.

### Case Study: Flowering Plants

To test for correlated rates of molecular and morphological evolution in empirical data, we applied the above five methods to large angiosperm data sets. Molecular data were obtained from the One Thousand Plant Transcriptomes Initiative (2019), hereafter ONEKP. This data set includes nucleotide sequences from 410 protein-coding nuclear genes from 1,124 green plants, glaucophytes, and red algae. For computational tractability, we analysed a subset of 111 angiosperm species and one gymnosperm outgroup (*Welwitschia mirabilis*), matching the angiosperm species in the morphological data set. We applied the data-partitioning scheme as outlined by ONEKP (2019) and estimated branch lengths on a fixed tree topology using maximum-likelihood analysis and Bayesian inference. The substitution model for each data subset was selected using ModelFinder in IQTREE2 (Kalyaanamoorthy et al. 2017).

The morphological data set was sourced from Schönenberger et al. (2020), representing a slightly expanded data set of 30 floral characters for 792 angiosperm species, initially published by (2017). These are discrete binary and multi-state morphological characters, including features such as the structural sex of flowers, ovary position, phyllotaxy, number of reproductive parts, and fusion of ovaries. We used a subset of 111 angiosperm species for our analyses and partitioned the data according to the number of character states (i.e., two-, three-, four-, and five-state data were treated as separate subsets). Branch lengths were estimated on a fixed tree topology using maximum-likelihood analysis and Bayesian inference, using the Mk+Γ model of evolution with correction for ascertainment bias. The model was selected using ModelFinder in IQ-TREE2.

We performed Bayesian phylogenetic analyses of the genomic data and floral characters with and without calibrations. In the former case, we implemented secondary calibrations on the ages of key angiosperm groups. The secondary calibrations were sourced from Ramírez-Barahona et al. (2020) and applied as normal priors on node ages (Ho and Phillips 2009). To estimate the posterior distribution, we sampled from five independent MCMC runs with chain lengths of either 50 or 70 million steps. Samples were drawn every 1000 steps. After checking for convergence and sufficient sampling in Tracer, we removed a burn-in fraction of between 40% and 60%, depending on the analysis, leaving a total of 170,000 sampled trees. Further details of the angiosperm case study, including settings and secondary calibrations, are available in the Supplementary Material.

We tested for correlations between rates of nuclear genomic evolution and floral character evolution using (*i*) root-to-tip distance correlations, (*ii*) independent sister-pairs contrasts, (*iii*) likelihood-based model selection, (*iv*) correlations of Bayesian branch rates, and (*v*) Bayesian model selection. We checked assumptions for each of these tests, as described in the Supplementary Material.

## Results

### Performance in Detecting Correlated Evolutionary Rates

Using data generated by simulation, we compared five methods for testing for correlations in evolutionary rates between molecular sequences and morphological characters. Correlations of branch rates from Bayesian relaxed-clock inference were able to detect correlated rates between molecular and morphological data under the widest range of simulation settings (Fig. 2 and 3). Under this method, the mean and median posterior branch rates were equally effective, with an average detection of correlated evolutionary rates of 85.8% and 85.6%, respectively. The performance of this method was closely followed by root-to-tip distance correlations (84.9%), Bayesian model selection (64.4%), and independent sister-pairs contrasts (48.2%) (Fig. 2; and 3). Likelihood-based model selection had very high detection of correlated evolutionary rates across scenarios (97.0% with AICc and 100% with BIC) but had an unacceptably high frequency of false positives.

**Figure 2.**
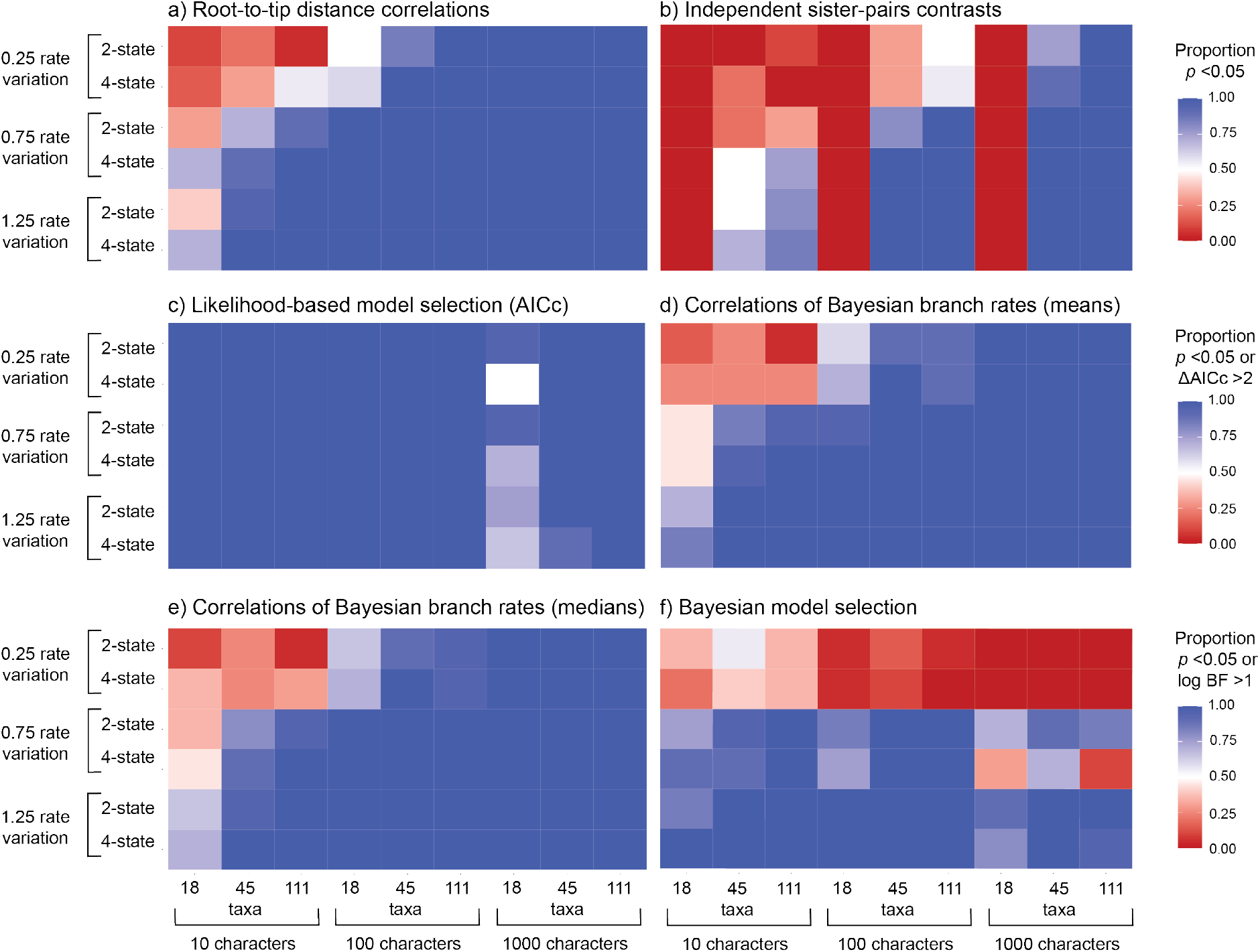
Heatmaps showing the performance of five approaches for testing for correlations between molecular and morphological evolutionary rates, for data produced by simulation with correlated evolutionary rates: a) root-to-tip distance correlations; b) independent sister-pairs contrasts; c) likelihood-based model selection using the corrected Akaike information criterion (AICc); correlations of Bayesian branch rates using d) mean posterior branch rates of branches or e) median posterior branch rates; and f) Bayesian model selection. In each panel, rows give results under six scenarios, representing combinations of three levels of among-lineage rate variation [0.25, 0.75, 1.25], and either two- or four-state morphological characters. In each panel, columns give results for data sets of various sizes, representing combinations of three numbers of morphological characters [10, 100, 1000] and three numbers of taxa [18, 45, 111]. For methods (a)–(b) and (d)–(e), colours indicate the proportion of 20 replicates for each setting that yielded a significant rate correlation (i.e., *p* < 0.05). For method (c), colours indicate the proportion of 20 replicates for each setting that yielded ΔAICc > 2, supporting a model of linked rates over a model of unlinked rates. For method (f), colours indicate the proportion of 20 replicates for each setting that yielded a log Bayes factor (BF) > 1.0 for a model of linked rates over a model of unlinked rates. For heatmaps of likelihood-based model selection using the Bayesian information criterion, see Supplementary Material.

**Figure 3.**
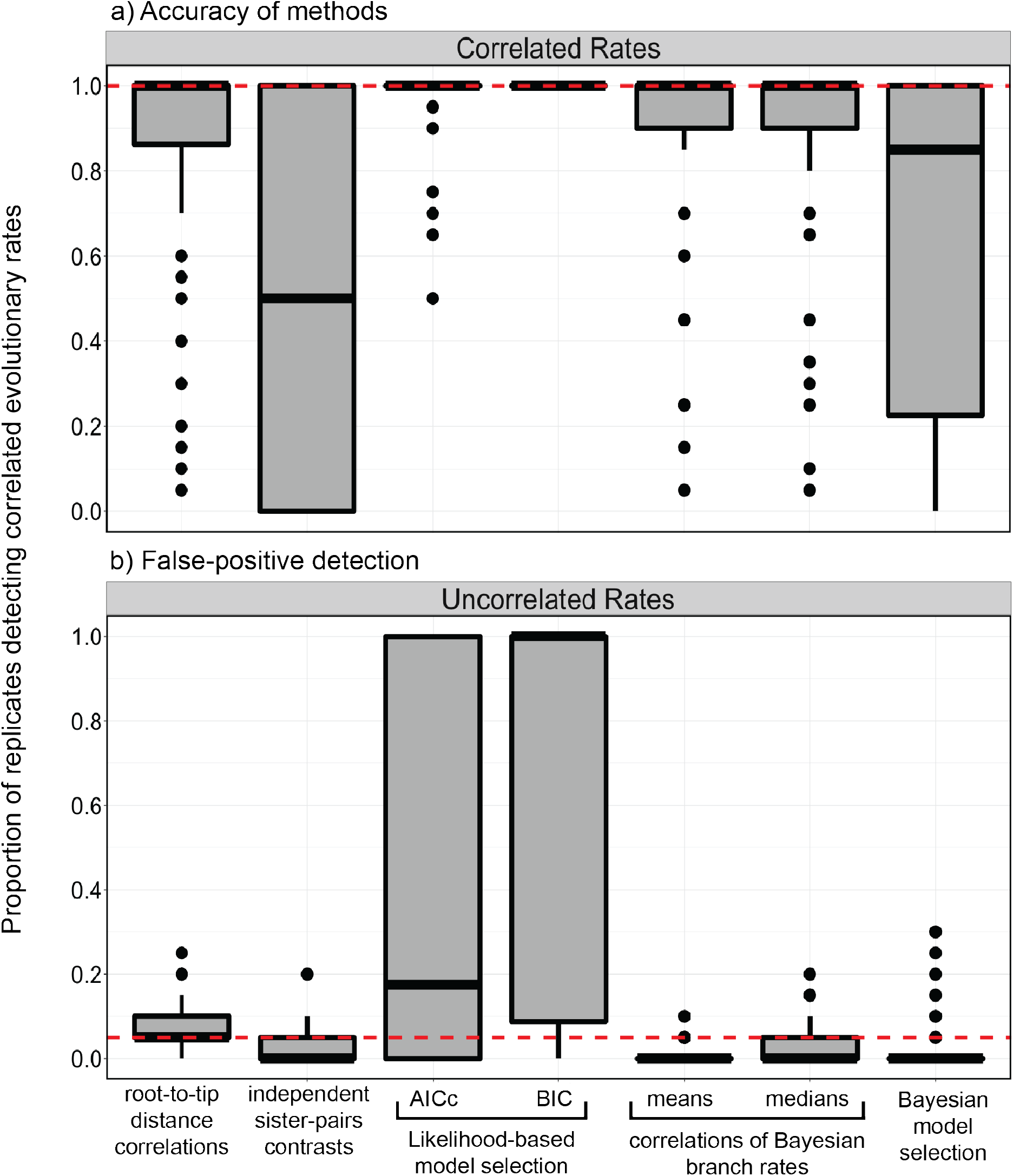
Boxplots showing the frequency of detecting correlated rates of evolution between simulated molecular and morphological data using different methods. A) Accuracy of the methods when analysing data simulated with correlated rates. The dashed horizontal line represents the ideal detection of correlated rates of evolution (100% of scenarios). B) Propensity of methods to detect correlations when analysing data simulated with uncorrelated rates (false positive detection). The dashed horizontal line represents the detection of correlated rates of evolution expected under frequentist statistics with a critical value of 0.05 (false positive rate of 5%).

When we analysed molecular and morphological data sets that had been generated without correlated rates, we found low false-positive rates when using correlations of Bayesian branch rates (1.20% and 2.96% for mean and median posterior branch rates, respectively), root-to-tip distance correlations (7.96%), independent sister-pairs contrasts (3.06%), and Bayesian model selection (3.06%) (Fig. 3 and 4). However, likelihood-based model selection yielded a high frequency of false positives when using either the AICc (47.3%) or BIC (69.3%).

**Figure 4.**
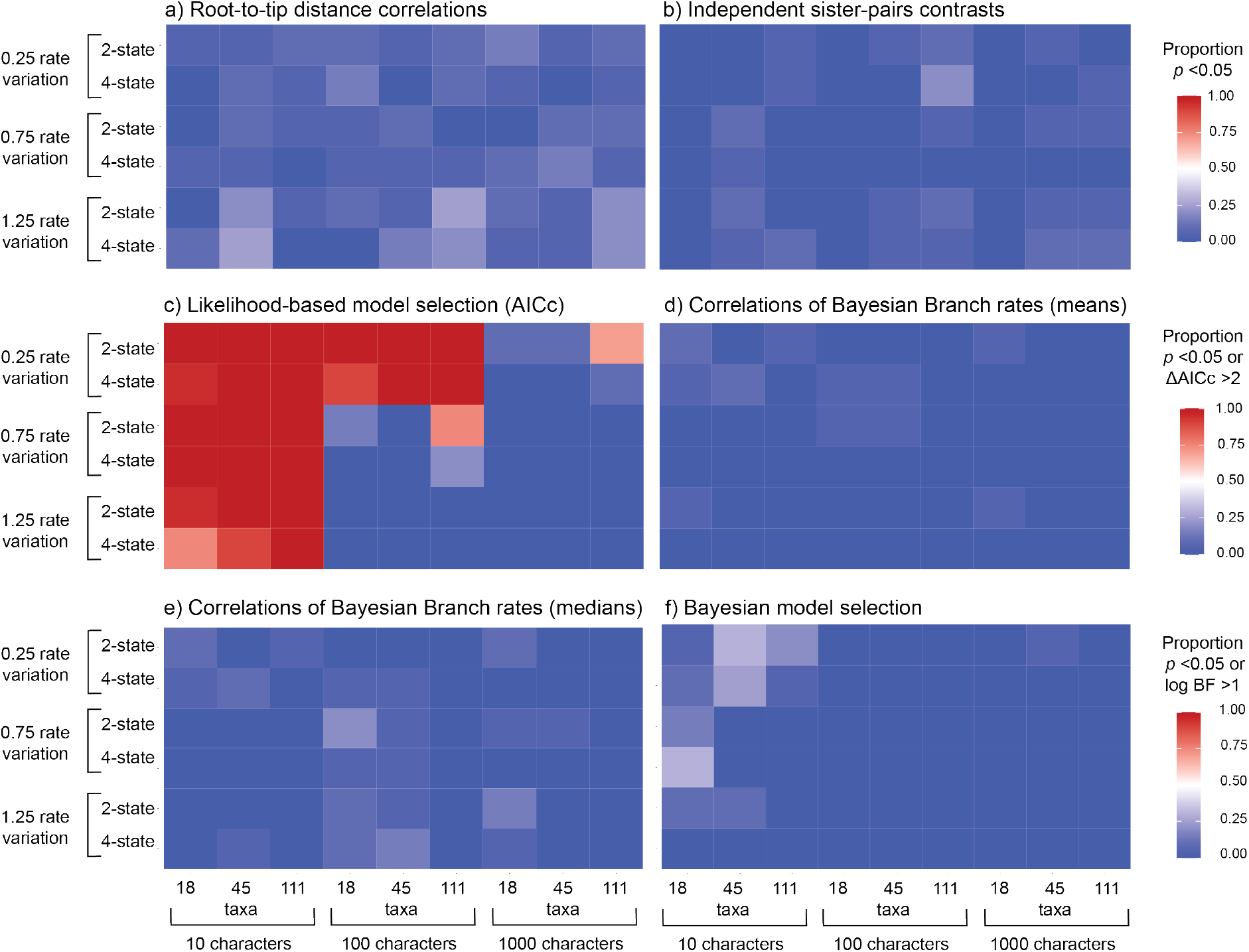
Heatmaps showing the performance of five approaches for testing for correlations between molecular and morphological evolutionary rates, for data produced by simulation without correlated evolutionary rates: a) root-to-tip distance correlations; b) independent sister-pairs contrasts; c) likelihood-based model selection using the corrected Akaike information criterion; correlations of Bayesian branch rates using d) mean posterior branch rates of branches or e) median posterior branch rates; and f) Bayesian model selection. In each panel, rows give results under six scenarios, representing combinations of three levels of among-lineage rate variation [0.25, 0.75, 1.25], and either two- or four-state morphological characters. In each panel, columns give results for data sets of various sizes, representing combinations of three numbers of morphological characters [10, 100, 1000] and three numbers of taxa [18, 45, 111]. For methods (a)–(b) and (d)–(e), colours indicate the proportion of 20 replicates for each setting that yielded a significant rate correlation (i.e., *p* < 0.05). For method (c), colours indicate the proportion of 20 replicates for each setting that yielded ΔAICc > 2, supporting a model of linked rates over a model of unlinked rates. For method (f), colours indicate the proportion of 20 replicates for each setting that yielded a log Bayes factor (BF) > 1.0 for a model of linked rates over a model of unlinked rates. For heatmaps of likelihood-based model selection using the Bayesian information criterion, see Supplementary Material.

### Impacts of Varying Simulation Conditions

We tested the effect of tree size, including 18, 45, and 111 taxa in the simulations, to evaluate its impact on the ability to detect correlated evolutionary rates. The number of taxa influenced the detection of correlations predictably (Fig. 5a), with performance increasing with tree size (57.7%, 81.1%, and 82.5% for tree sizes of 18, 45, and 111 taxa, respectively). Analyses of the 18 taxon-set performed poorly when there were only 10 morphological characters (Fig. 2). This was especially apparent for independent sister-pairs contrasts, where analyses of 18-taxon data sets failed to detect rate correlations regardless of the degree of among-lineage rate variation, number of character states, and numbers of morphological characters.

**Figure 5.**
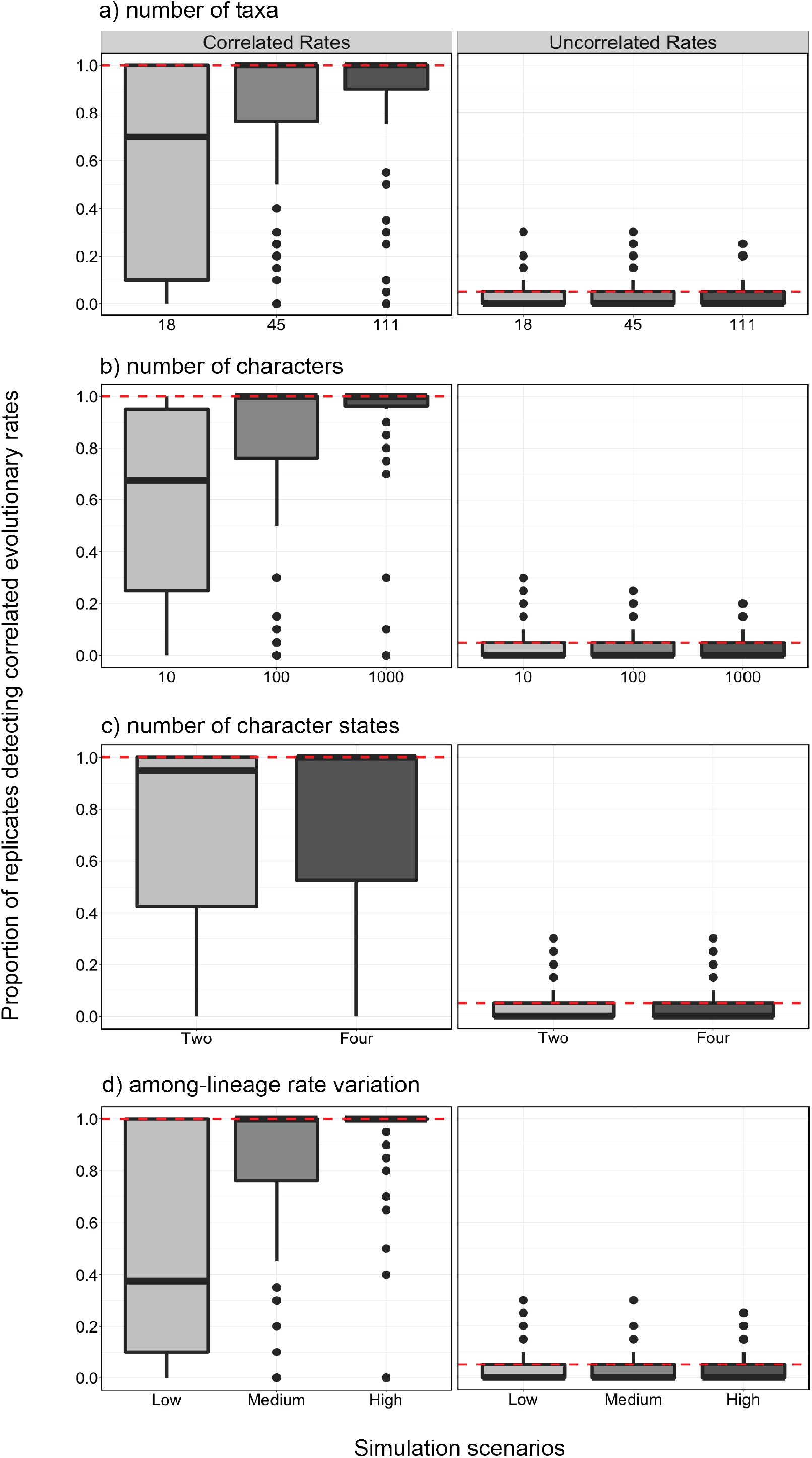
Boxplots showing the frequency of detecting correlated rates of evolution between simulated molecular and morphological data using different settings. The left panels show the performance of methods when data were simulated with correlated rates, with dashed horizontal lines representing the ideal detection of correlated rates of evolution (100% of scenarios). The right panels show results when simulated with uncorrelated rates, with dashed horizontal lines representing the detection of correlated rates of evolution expected under frequentist statistics with a critical value of 0.05 (false positive rate of 5%). The different settings used were: a) three tree sizes, b) three sizes of morphological character matrices, c) two numbers of possible morphological character states, and d) three levels of among-lineage rate variation. The results were pooled across all methods except for likelihood-based model selection, since this method had such a high rate of false positives and would unreasonably skew the detection of correlations. Boxplots calculated with the results from likelihood-based model selection can be found in the Supplementary Material.

We varied the number of morphological characters [10, 100, 1000] to evaluate their impact on the ability of the five methods to detect correlated evolutionary rates. We found that the average detection of positive correlations increased from 57.6%, 80.6%, to 83.2% for data sets with 10, 100, and 1000 morphological characters, respectively (Fig. 2 and 5b). We found that the effect depended on the amount of among-lineage rate variation; where branch rates had a standard deviation of at least 0.75, the four best approaches were generally able to detect correlations with any number of morphological characters (Fig. 2). However, when there was a low degree of among-lineage rate variation, rate correlation could not be detected when there were only 10 morphological characters. Furthermore, likelihood-based model selection detected a high rate of false positives, but this was mitigated when there were either 100 or 1000 morphological characters and moderate or high among-lineage rate variation (Fig. 4c and Supplementary Material).

The number of character states for the morphological data had a minor impact on detection of correlations. Generally, correlations were more frequently detected when the morphological data comprised four-state characters, with a positive detection of 75.1% compared with 72.4% when the data comprised two-state characters (Fig. 2 and 5c), although this effect was negligible when there were greater than 10 morphological characters and 18 taxa in the data set (Fig. 3).

Across the four most powerful and accurate methods, the most important factor for detection of correlated evolutionary rates was among-lineage rate variation (Fig. 2 and 5d). When we implemented low among-lineage rate variation in the data sets, correlated rates could only be detected across 50.2% of replicates, whereas medium and high levels increased detection to 81.9% and 89.2%, respectively. Where there was low among-lineage rate variation, with branch rates having a standard deviation of 0.25, the four best approaches were generally unable to detect correlations without sampling at least 100 morphological characters (Fig. 2). This was especially true for Bayesian model selection, which could not detect correlated rates of evolution at the lowest level of among-lineage rate variation.

### Case Study: Flowering Plants

In our analyses of genomic DNA and floral characters in angiosperms, we found that two of the five methods, root-to-tip distance correlations and likelihood-based model selection, detected a correlation in evolutionary rates. We found evidence of a correlation in our analysis of root-to-tip distances (*p* ≈ 0; Fig. 6a). Outliers (data points outside 1.5 times the interquartile range) were excluded from the permutation test, but root-to-tip distances were significantly correlated both before and after removal of outliers (see Supplementary Material). These outliers included the branch leading to the sister taxon to all remaining angiosperms, *Amborella trichopoda*, which had a low morphological evolutionary rate of 2.38×10^−6^ changes/character/Myr. Other ANA-grade angiosperms, such as *Austrobaileya scandens* and *Illicium floridanum*, similarly had low rates of morphological change and were removed from the test. Likelihood-based model selection also yielded strong support for linking branch lengths between nuclear sequences and floral characters, when using both AICc (ΔAICc = 203.4) and BIC scores (ΔBIC = 2504.9).

**Figure 6.**
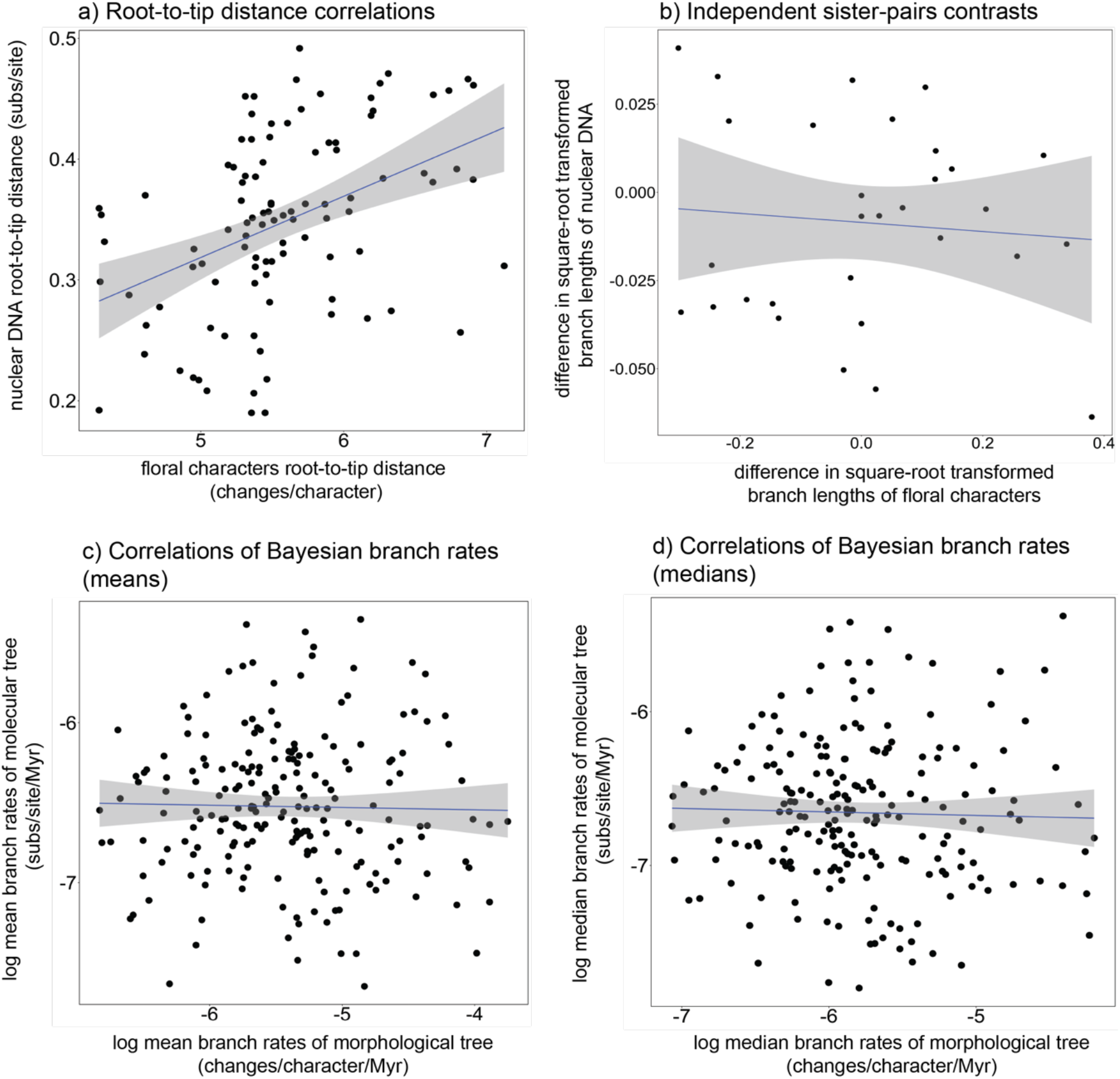
Comparisons between evolutionary rates of nuclear genomic DNA and floral characters inferred from angiosperms. The methods used to test for correlated rates of evolution are a) root-to-tip distance correlation; b) independent-sister pairs contrasts; c) correlations of Bayesian mean posterior branch rates; and d) correlations of Bayesian median posterior branch rates. The plot for each comparison has been fit with a linear model, which is displayed along with the 95% confidence interval.

We found no evidence of correlated molecular and morphological rates when we analysed the data using independent sister-pairs contrasts (*r*_*s*_ = 0.0162, *p* = 0.466; Fig. 6b), correlations of Bayesian mean posterior branch rates (*r* = –0.0209, *p* = 0.617; Fig. 6c), or correlations of Bayesian median posterior branch rates (*r*_*s*_ = –0.0685, *p* = 0.837; Fig. 6d). The Bayes factor gave very strong support to unlinking the clock models between nuclear and floral characters, with an average log Bayes factor of 51.4. Our Bayesian approaches failed to detect a correlation regardless of whether fossil calibrations were included or not (see Supplementary Material for the results of analyses without fossil calibrations).

## Discussion

We have shown through a comprehensive simulation study that correlated rates of evolution between molecular sequences and morphological characters can be detected under a variety of circumstances. The best-performing method was correlations of Bayesian branch rates, followed by root-to-tip distances, Bayesian model selection, independent sister-pairs contrasts, and lastly likelihood-based model selection. However, when taking computational burden into account, testing for correlations using root-to-tip distances is the most efficient method. Overall, methods had more power when the data had a high degree of among-lineage rate variation, and when at least 45 taxa or 100 morphological characters were sampled. When we applied these methods to an angiosperm data set, we found limited evidence for coupled evolutionary rates when analysing nuclear DNA and floral characters. The estimation of root-to-tip distances might have been misled by missing character data in the floral trait data set. However, missing data are often unavoidable in morphological data sets, due to inapplicable characters, i.e., characters that are not common across species, and difficulties in accessing suitable samples (Scholtz 2010; Wanninger 2015). Although our simulations used evolutionary parameters that were empirically informed, they still represented an idealized form of the evolutionary process and yielded complete data sets. Below we discuss the results and implications of the simulation study before returning to the case study of angiosperms.

### Detecting Correlations Between Rates of Molecular and Morphological Evolution

Our simulation study has provided detailed evaluations of five methods of testing for correlations between rates of molecular and morphological evolution. We found that the three methods that used inferences from maximum-likelihood phylogenetic analysis required at least 100 morphological characters for accurate detection of rate correlations. We found that correlations of root-to-tip distances performed well, with a low rate of false positives. While statistical analyses of root-to-tip distances are hindered by the non-independence of the data points (Rambaut et al. 2016), appropriate *p*-values can be computed using a permutation test (Higgins 2004; Garren 2019). Analysis using independent sister-pair contrasts was able to detect rate correlations less frequently than the other methods that we evaluated, and this is likely to be due to the reduced number of data points that are sampled. For instance, for the tree including 111 species, root-to-tip distance correlations are based on 111 data points, whereas independent sister-pairs contrasts use only 35 data points.

When we used likelihood-based model selection to compare models with proportionate (linked) versus unlinked branch lengths, we consistently found support for the proportionate model even for data that had been generated by simulation with uncorrelated rates. However, this was probably because the proportionate model captures a substantial amount of variation while bringing only a modest increase in the number of parameters (Duchêne et al. 2020). The proportionate model was favoured under almost all simulation settings, except when there was a large number of morphological characters. Of the two information criteria that were employed, model selection using the AICc yielded fewer false positives, since it penalizes model size less harshly than the BIC (Duchêne et al. 2020).

The two Bayesian methods of testing for rate correlations showed highly contrasting performance in our simulation study. We found that correlations could be detected in our analyses of Bayesian branch rates even under less informative settings, such as when there were only 10 morphological characters. This strong performance was unexpected, given the typically large uncertainty in estimates of branch rates (e.g., Ho et al. 2005; Drummond et al. 2006), but can perhaps be attributed to the large number of data points sampled by the test (one comparison per branch). Compared with the likelihood-based approach, Bayesian model selection performed well when detecting correlated evolutionary rates, except when there was low among-lineage rate variation. It may be useful to compare the performance of other methods of marginal-likelihood estimation, such as path sampling or nested sampling (Skilling 2006; Russel et al. 2019).

Further evaluations of methods that test for correlations between molecular and morphological rates of evolution will be valuable, given that the dynamics of morphological evolution and the relationship to molecular evolution remain poorly understood (Lee and Palci 2015). In our study, we have not considered processes ‘external’ to coding in DNA sequences, such as phenotypic plasticity and epigenetics, but these may shape adaptation and phenotypic changes over time (West-Eberhard 1989; Nylin and Wahlberg 2008). Furthermore, there is a lack of congruence between phylogenies inferred from different types of biological data (Oyston et al.), possibly due to the limited size of morphological data sets or the effect of homoplasy (Keating et al. 2020). However, this might not be pertinent at higher taxonomic levels (Jablonski and Finarelli 2009), where diagnostic characters tend to be more informative and can carry strong phylogenetic signal.

By testing for correlations between molecular and morphological evolutionary rates, we can better understand the dynamics of the ‘morphological clock’. Whilst a broad link between molecular and morphological change is expected (Simpson 1953), the existence of a morphological clock has so far been rejected (Beck and Lee 2014; Lee and Palci 2015; O’Reilly et al. 2015; Lee 2016; Tarasov 2019). This is reinforced by the apparent lack of a ‘common mechanism’ governing the evolution of morphological characters, with the pattern of among-character rate variation differing across branches (Goloboff et al. 2019).

The performance of methods is likely to be worse in analyses of real data sets compared to simulated data sets, because of the complexities of the evolutionary process in reality and because of the shortcomings of our evolutionary models, particularly models of morphological evolution. Estimating morphological rates of evolution is fraught with uncertainty, and the distribution of rates across taxa and over time is largely undescribed (Simpson 1944; Schopf 1984). Unlike molecular data, the collection of morphological data is ‘infinitely extensible’; there is no upper boundary on the total number of characters and states that can be considered (Oyston et al.), because there are no objectively defined categories such as the 20 amino acids or four nucleotides found in molecular data (Dávalos et al. 2014; Lee and Palci 2015; Barba-Montoya et al. 2021). The morphological characters that are selected for phylogenetic inference are usually chosen for their diagnostic utility, so invariant and rapidly evolving characters are typically excluded (Lewis 2001; Wright and Hillis 2014).

Previous work has shown that Bayesian and maximum-parsimony phylogenetic analyses of morphological data have greater accuracy for data that have been generated under stochastic processes rather than being subject to selection (Keating et al. 2020). This indicates that at a macroevolutionary scale, the dynamics of morphological evolution may deviate from the Mk model, which is a simplified version of the general multiple-rate asymmetrical Mk model, originally introduced for morphological data (Pagel 1994; Goloboff et al. 2019; Keating et al. 2020). Although the inadequacy of the Mk model is often assumed to hamper phylogenetic inference using morphological characters, it might not be a substantial problem unless homoplasy is particularly extensive (Jablonski and Finarelli 2009) or when rates are extremely high (Reyes et al. 2018; Klopfstein et al. 2019; Simões et al. 2022a). Apart from these cases, Bayesian inference using the Mk model seems to be relatively robust and can accurately infer topologies and branch lengths under a broad range of conditions (Klopfstein et al. 2019).

### Evolutionary Rates in Angiosperms

Our analysis of data from 111 angiosperms failed to detect a correlation between the evolutionary rates of nuclear DNA and floral traits. The 30-character floral data set that we analysed might not have been sufficiently informative, so our results will require confirmation using larger data sets comprising at least 100 characters. Additionally, the floral data set that we examined excluded hypervariable characters, such as floral colour. The resulting characters in the floral data set were all slowly evolving, at rates below 0.006 changes/Myr. These low rates have been described as ‘optimal’ for phylogenetic inference (Klopfstein et al. 2019), and were likewise suited to the primary goal of ancestral state reconstruction for which this data set was assembled (Sauquet et al. 2017). However, simulating data sets with a broader diversity of rates, including both higher and lower ones, would be useful. Also, incorporating missing data in the simulations would improve the realism of the data sets and allow evaluation of the impacts of missing data on detecting correlations in evolutionary rates.

The molecular data set used to test for correlations between rates of floral and sequence evolution included 410 protein-coding, single-copy nuclear genes, obtained by sequencing the vegetative tissue transcriptomes of plant species (ONEKP 2019). These 410 protein-coding genes likely control phenotypic expression across a broad range of characters. However, the set of 30 curated floral traits only represents a small proportion of the total phenotypic traits of each flowering plant species. Assessing a larger number of morphological traits will not only lend more power to the analyses but will also provide a more accurate reflection of the overall rate of morphological trait evolution. Such a data set is not yet available for flowering plants across a broad phylogenetic sample because of the considerable work required in its assembly. Studies could possibly be done with resources such as the global ‘TRY’ plant database (Kattge et al. 2020), which is composed mostly of vegetative traits. Generally, floral traits are under intense sexual selection (Barrett 2010), which might influence the detection of correlated rates. Indeed, Barraclough and Savolainen (2001) found a very strong correlation between the evolution of molecular sequences and floral traits, but a weak correlation when analysing vegetative traits.

Overall, the result might correctly reflect a more general uncoupling of molecular and morphological rates in angiosperms. A decoupling of evolutionary rates between the nuclear genome and floral characters suggests a departure from a model of gradual morphological change, i.e., that morphological evolution is not proportional to time (Halliday et al. 2019). This may be because the floral characters exhibit high heterogeneity and deviation from clocklike evolution. Indeed, from the Bayesian relaxed-clock analysis, the floral characters exhibited a coefficient of variation of branch rates of 1.53 (95% credible interval 0.896–2.24) whereas the genomic DNA had a coefficient of variation of 1.35 (95% credible interval 1.27– 1.52). Pulses of morphological change have occurred throughout plant evolution, possibly at speciation events (Eldredge and Gould 1972), with notable episodes corresponding to the introduction of vascular plants in the Devonian and the diversification of angiosperms in the Late Cretaceous (Leslie et al. 2021).

A lack of an association between rates of floral character and molecular evolution would also be consistent with floral evolution being driven by changes at specific loci (Kimura 1968; Barrier et al. 2001; Davies and Savolainen 2006; Duret 2008; Gaut et al. 2011). The mutations that produce phenotypic change might occur largely in adaptive and regulatory genes, while many genomic mutations are neutral in their impact on fitness (Kimura 1968, 1983). Indeed, a large proportion of the morphological diversity amongst flowering plants can be attributed to specialized interactions between angiosperms and their insect pollinators (Darwin 1862; Friis et al. 2006; Benton et al. 2021; Asar et al. 2022). Furthermore, in our study, we were limited to examining evolutionary change in protein-coding genes, which only included the first two sites of each codon. Testing for correlations separately using rates of nonsynonymous and synonymous substitution will allow further insights into the relative importance of selection and drift (Barrier et al. 2001).

### Concluding Remarks

We have shown that correlations between molecular and morphological evolutionary rates can be detected under the conditions explored in our simulation study. However, the complexities of how morphological evolution proceeds, and whether this is effectively described by current evolutionary models and approaches, will ultimately determine whether the rates of morphological character evolution and their correlates can be accurately reconstructed in practice. While we did not find evidence of correlated evolutionary rates between angiosperm genomic DNA and floral characters, the question of whether the rates of genotypic and phenotypic evolution are correlated in angiosperms should be addressed with a larger morphological data set.

Our study has implications for combined analyses of molecular and morphological data, where branch lengths between data sets are often linked as a default approach (Nylander et al. 2004; O’Reilly et al. 2015). The results of our simulation study lead us to suggest that future studies should use morphological character matrices of at least 100 characters; this would allow for partitioning of the morphological data set, which has been demonstrated to improve the precision of divergence date estimates and accuracy of branch-length estimates (Lee 2016; Caldas and Schrago 2019; Neumann et al. 2021). Moreover, increasing the size of the morphological data set can minimize the impacts of character correlation (Guillerme and Brazeau 2018; Simões et al. 2022a). This work should not only be extended to larger data sets, but should also span across the Tree of Life, to help elucidate the processes that drive macroevolutionary change. Furthermore, these methods are not restricted to analyses of molecular and morphological evolution, but can also be used to test for correlations in rates between symbionts and their hosts or between organellar and nuclear genomes in plants.

## Supplementary Materials

Supplementary material is available online. All text, files and code are available at Dryad [X].

## Acknowledgements

Y.A. acknowledges the support provided by the Australian Government’s Research Training Program award. This work was funded by the Australian Research Council (DP220103265). The authors acknowledge the Sydney Informatics Hub and the high-performance computing facility, Artemis, at the University of Sydney for providing computing resources.

## Notes

### Competing Interest Statement

The authors have declared no competing interest.

